# Using Earth Mover’s Distance for Viral Outbreak Investigations

**DOI:** 10.1101/628859

**Authors:** Andrew Melnyk, Sergey Knyazev, Fredrik Vannberg, Leonid Bunimovich, Pavel Skums, Alex Zelikovsky

## Abstract

RNA viruses mutate at extremely high rates forming an intra-host viral population of closely related variants (or quasi-species) [4]. High variability of Human Immunodeficiency Virus (HIV) and Hepatitis C virus (HCV) making them particularly dangerous by allowing them to evade the host’s immune system. HIV and HCV outbreaks pose a significant problem for public health for solving which it is critical to infer transmission clusters, i.e., to decide whether two viral samples belong to the same outbreak. Initial approach [10] was based on estimating relatedness between two samples as the distance between consensuses of the corresponding viral populations. The distance between closest pair of representatives from two populations, *MinDist*, has been shown to be significantly more accurate [2]. Unfortunately, *MinDist* computation requires a cumbersome RNA-seq data assembly and identification of all viral sequences from a given project. We present a novel approach that allows to bypass read assembly and estimate the distance between viral samples based on k-mer (i.e. a substring of length k) distribution in RNA-seq reads. The experimental validation using sequencing data from HCV outbreaks shows that the proposed algorithms can successfully identify genetic relatedness between viral populations, infer transmission clusters and outbreak sources, as well decide whether the primary spreader is present in the sequenced outbreak sample.

## 1 Introduction

RNA viruses mutate at extremely high rates forming an intra-host viral population of closely related variants (or quasi-species). High variability of Human Immunodeficiency Virus (HIV) and Hepatitis C virus (HCV) making them particularly dangerous by allowing them to evade the host’s immune system. HIV and HCV outbreaks pose a significant problem for public health for solving which it is critical to infer transmission clusters, i.e., to decide whether two viral samples belong to the same outbreak.

The progress of sequencing technologies made possible to identify and sample intra-host viral populations at great depth. Consequently, a contribution of sequencing technologies to molecular surveillance of viral disease epidemic spread becomes more and more substantial. Genome sequencing of viral populations reveals similarities between samples, allows measuring viral genetic distance, and facilitates outbreak identification and isolation. Computational methods can be used to infer transmission characteristics from sequencing data. The popular high-throughput sequencing (HTS) technology MiSeq is used to sequence viral samples and detect rare viral mutations. Short MiSeq reads complicates alignment and assembly of rapidly mutating RNA viruses such as HCV and HIV and skipping these error prone and time steps become highly attractive. In this paper, we propose to apply the alignment- and assembly-free *k*-mer strategy to viral sequencing data. This strategy has been initially introduced for analyzing HTS data in metagenomic studies where reads come from multiple related and unrelated genomes (see [6]) as well as for RNA-seq quantification [1].

Indeed, it is relatively fast and easy to extract k-mers from reads, and so the complexity of viral distance measurement shifts from assembly to a comparison of k-mer sets or distributions. Following [6], we build a De Bruijn graph for each sample and then calculate an Earth Mover’s Distance (EMD) between two *k*-mer distributions.

We applied the *k*-mer strategy to the following well-known epidemiological tasks (1-4).

T1. **Identification of primary spreader:**

**Given:** A set of hosts from the same viral outbreak
**Find:** The host which is the primary spreader of infection

T2. **Presence of primary spreader in the outbreak:**

**Given:** A set of hosts from the same viral outbreak
**Find:** Decide whether the primary spreader is among the hosts.

T3. **Detection of transmission clusters:**

**Given:** A set of related and unrelated hosts
**Find:** The transmission clusters corresponding to the separate outbreaks

T4. **Inference of transmission direction problem:**

**Given:** A pair of hosts, belonging to the same viral outbreak
**Find:** The direction of transmission between hosts

Identifying the main spreader of an outbreak is a crucial epidemiological task (T1) which helps intermit outbreak spreading. But the source can frequently be absent in collected data, so another critical task is to decide whether the source is among collected samples (T2). Knowing boundaries of an outbreak can help to measure infection spread, an efficiency of interventions into the outbreak, and be focused on the outbreak’s participants. So the next important task is discovering the boundaries of an outbreak (T3). The last primary task is decide who infected whom which necessary for finding the transmission chains (T4).

The experimental validation using sequencing data from HCV outbreaks shows that the porposed *k*-mer strategy can successfully identify genetic relatedness between viral populations, infer transmission clusters, outbreak sources, and transmission direction as well decide whether the primary spreader is present in the sequenced outbreak sample.

## Methods

Our algorithms are based on finding the distance between population using *Earth Movers’ Distance (EMD)* between corresponding distributions of k-mers and finding centers of a set of hosts (i.e., intra-host populations) represented by mean *k*-mer distributions. The general pipeline of the algorithm (see Figure 1) includes obtaining k-mer distributions from NGS reads for corresponding hosts and computing EMD between them. We first describe how we find distances between k-mers and then describe how we find distance between samples.

**Fig. 1:**
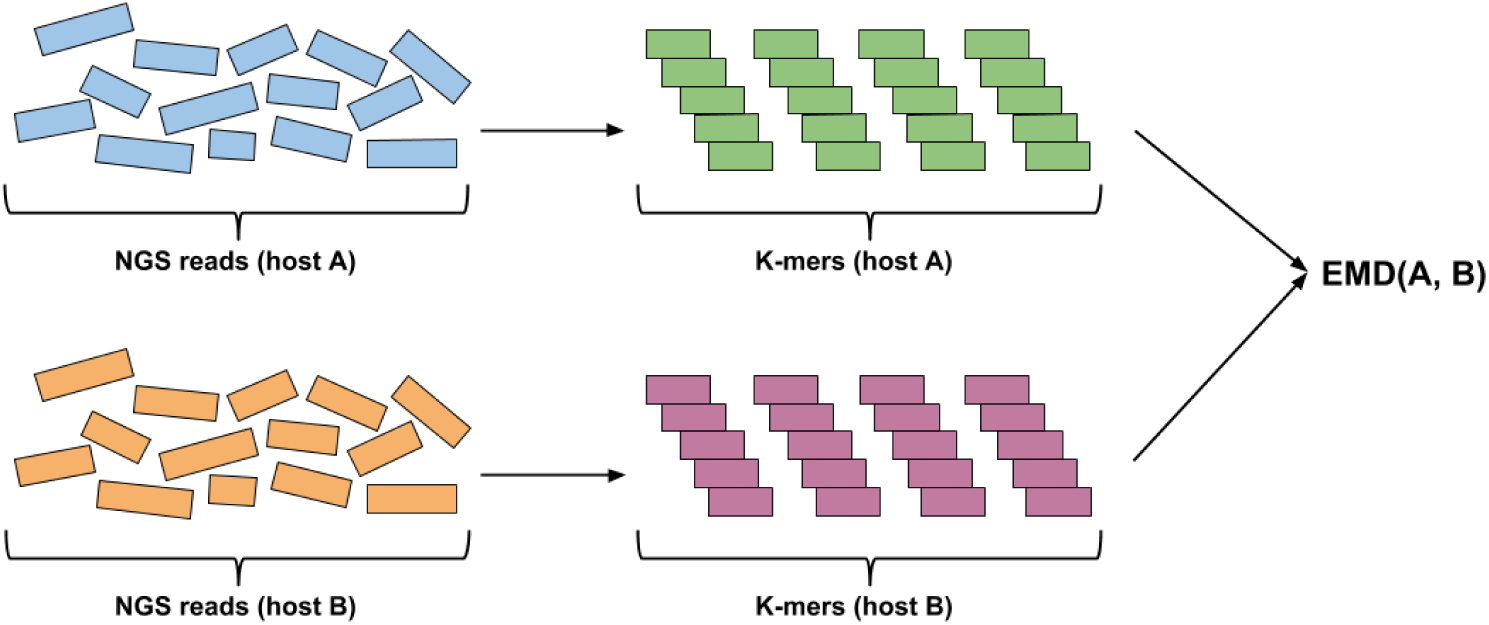
Algorithm pipeline. K-mer distributions for hosts, that need to be compared, are obtained from NGS reads. Then, EMD is computed using k-mer distributions.

### Finding distances between k-mers in the De Bruijn graph

*k-mer* refers to a substring of length *k*. In our work, we use *De Bruijn graph* to calculate distance between k-mers. De Bruijn graph is the graph, that is constructed so that vertices represent every string over a finite alphabet of length *l*, and edges are added between vertices that have overlap of *l* − 1.

Once De Bruijn graph is constructed, distance between k-mers can be calculated as a length of shortest path between corresponding vertices using *breadth-first search* algorithm. In our algorithms, obtained graph is converted to undirected before shortest path computation.

### Finding EMD between viral samples

Viral populations can be compared by comparing the corresponding k-mer distributions using EMD. First, K-mer distributions are obtained for each sample, so that they contain all k-mers and normalized frequencies.

EMD is a method, that allows to evaluate dissimilarity between two multi-dimensional distributions in some feature space where a distance measure between single features (*ground distance*) is given [8]. Distributions can be represented as *signatures* – sets of clusters, so that each cluster is represented by its mean and by the fraction of distribution that belongs to that cluster. Computation of EMD is based on solving the *transportation problem*, which can be formulated as following: for several suppliers, each with a given amount of goods, several consumers, each with limited capacity, and a cost of transporting a single unit of goods between each supplier-consumer pair, find a least-expensive flow of goods from the suppliers to the consumers that satisfies the consumers’ demand. EMD is calculated as the following *EMD* 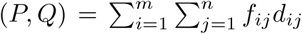 where *f*_*ij*_ is the minimum-cost flow between supplier *i* and consumer *j*, and *d*_*ij*_ is the distance be-tween *i* and *j*. It should also be noted that EMD is usually normalized by the total flow, but we perform normalization of frequencies in k-mer distributions before EMD computation, which results in total flow always being equal to 1.

#### K-EMD distance computation

**Input:** Sets of sequencing reads from hosts *A* and *B* (*S*_*A*_ and *S*_*B*_).

**Output:** k-EMD distance between *A* and *B*.

1. Produce k-mers from *S*_*A*_ and *S*_*B*_:

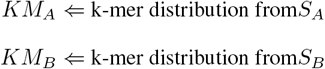
2. Initialze distance matrix *D*(*A, B*): for any pair of k-mers *x* ∈ *KM*_*A*_ and *y* ∈ *KM*_*B*_, find *dist*(*A, B*) in De Bruijn graph;
3. Compute *EMD*(*KM*_*A*_, *KM*_*B*_, *D*(*A, B*)).

Constructing of the De Bruijn graph between two sequences *GACTACTACTGT* and *GACTAGTACTGT* is shown on Figure 2. Figure 3 describes an example of k-EMD distance computation. After k-mer distributions are generated for input sequences, EMD is computed as the work 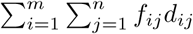, where *f*_*ij*_ is the flow between histogram(k-mer distribution) elements *i* and *j* and *d*_*ij*_ is the corresponding distance between k-mers, which is obtained from De Bruijn graph (Figure 2). This way, *EMD* = 1.625.

**Fig. 2:**
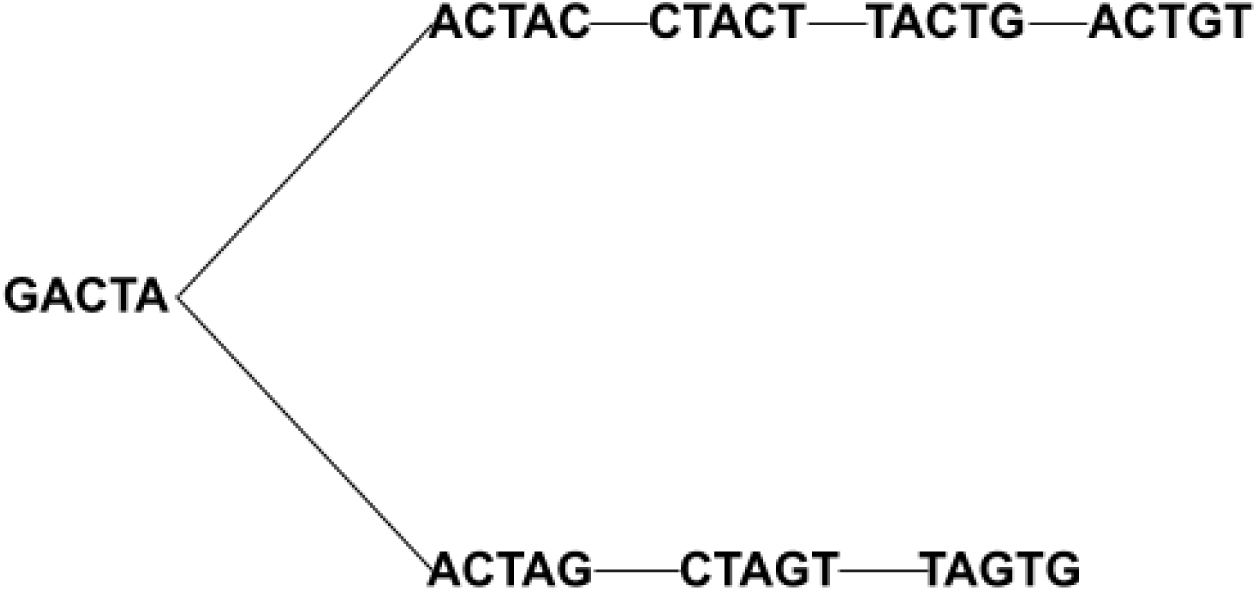
De Bruijn graph for 5-mers, obtained from sequences *GACTACTACTGT* and *GACTAGTACTGT*.

**Fig. 3:**
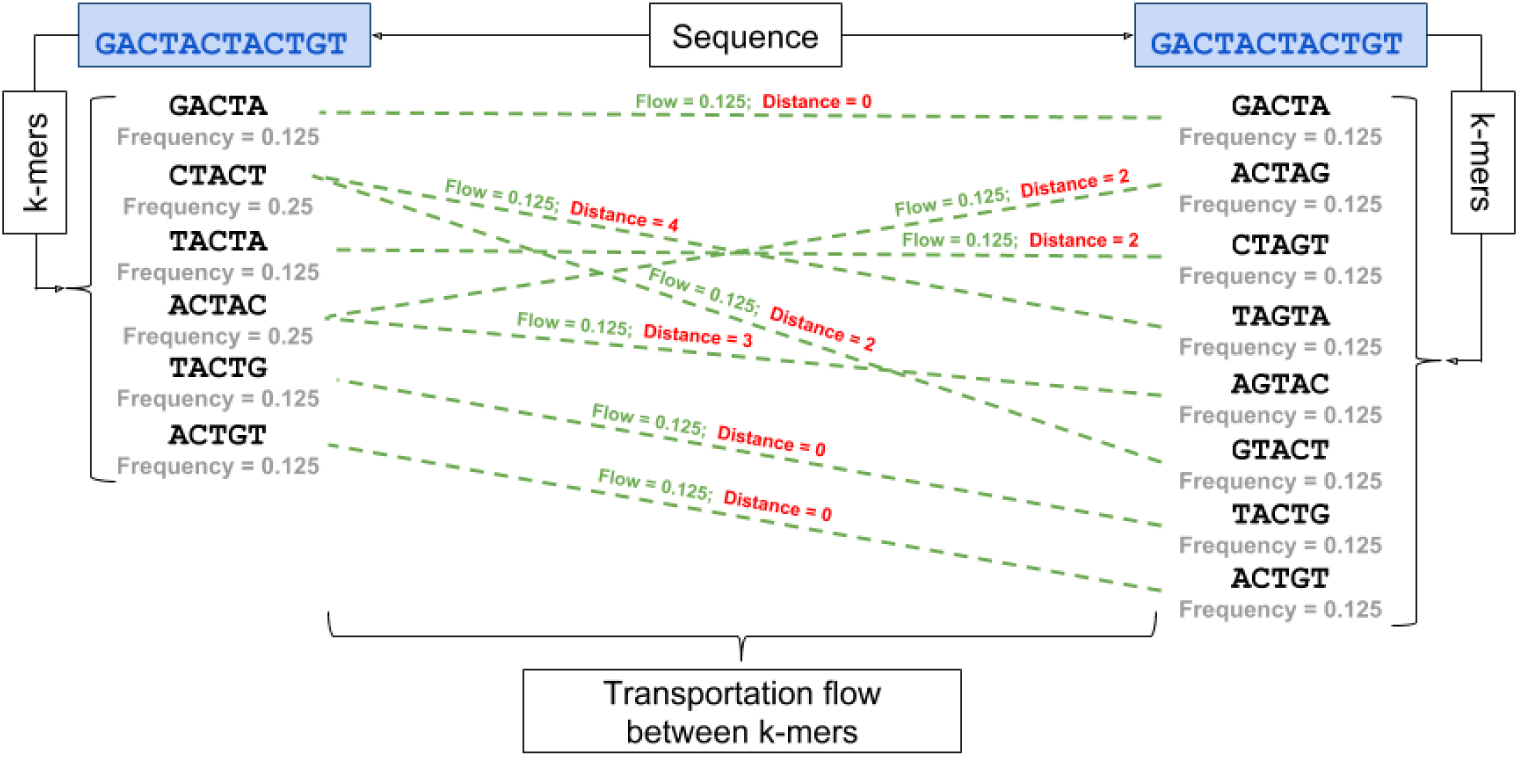
Finding EMD distance between *k*-mers of sequences *GACTACTACTGT* and *GACTAGTACTGT*. The *k*-mer distributions are on the left and right sides. Dashed lines represent transportation flow between k-mers; corresponding flow values are shown in green. Red values on top of the lines represent distance between corresponding k-mers in the De Bruijn graph.

### Mean distribution

Representing samples as k-mer distributions allows to estimate the center from a group of samples by introducing an arbitrary host, which can either be an *arithmetic mean, quadratic mean* or a *maximum mean*.

**–** The **arithmetic mean** *k*-mer distribution can be calculated by constructing a union 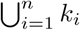, which contains all k-mers from a given set of samples and their respective frequencies 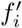, obtained as mean frequencies for given k-mer in analyzed set of samples 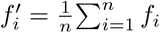.
**–** For the **quadratic mean** *k*-mer distribution, 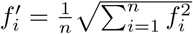.
**–** The **maximum mean** *k*-mer distribution is obtained by considering the maximum observed frequency for each k-mer 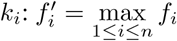.

### Hierarchical clustering

To test hierarchical clustering, *single-linkage* algorithm was used. This method evaluates the similarity of two clusters based on their most similar members [7] and groups clusters in bottom-up order until certain termination condition is satisfied. In our algorithm, we use a distance criteria, so clusters are merged until distance between them exceeds a pre-defined distance threshold, which represents EMD between two closest unrelated samples in the dataset. This way, we obtain a partition, where some of the related hosts remain in different clusters. At this point, we proceed to the second stage of the algorithm, that allows to improve the clustering quality by merging the clusters, that contain related hosts by performing the following steps:

1. For each cluster, obtained from hierarchical clustering, compute center as the mean of all hosts within the cluster;
2. For each center, obtained at the previous step:
  **–** Find distances to the furthest in-cluster host and closest host, that belongs to the different cluster;
  **–** If for cluster *A* there exists an ‘overlap’ (there is a host from cluster *B*, that is closer to the center than the furthest host, belonging to the same cluster (*A*)), merge *A* and *B*

Example of the algorithm is demonstrated in Figure 4, where related clusters, belonging to outbreak *AI* were merged after detecting an overlapping host *AI*004.

**Fig. 4:**
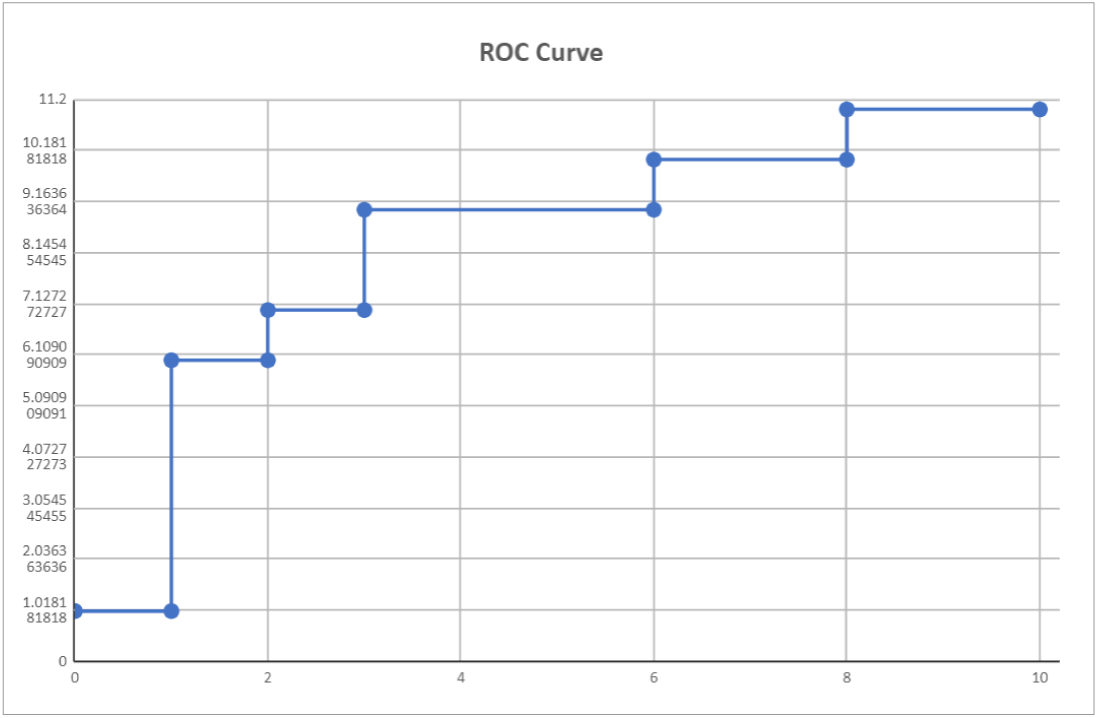
ROC curve for outbreak sources detection.

**Fig. 5:**
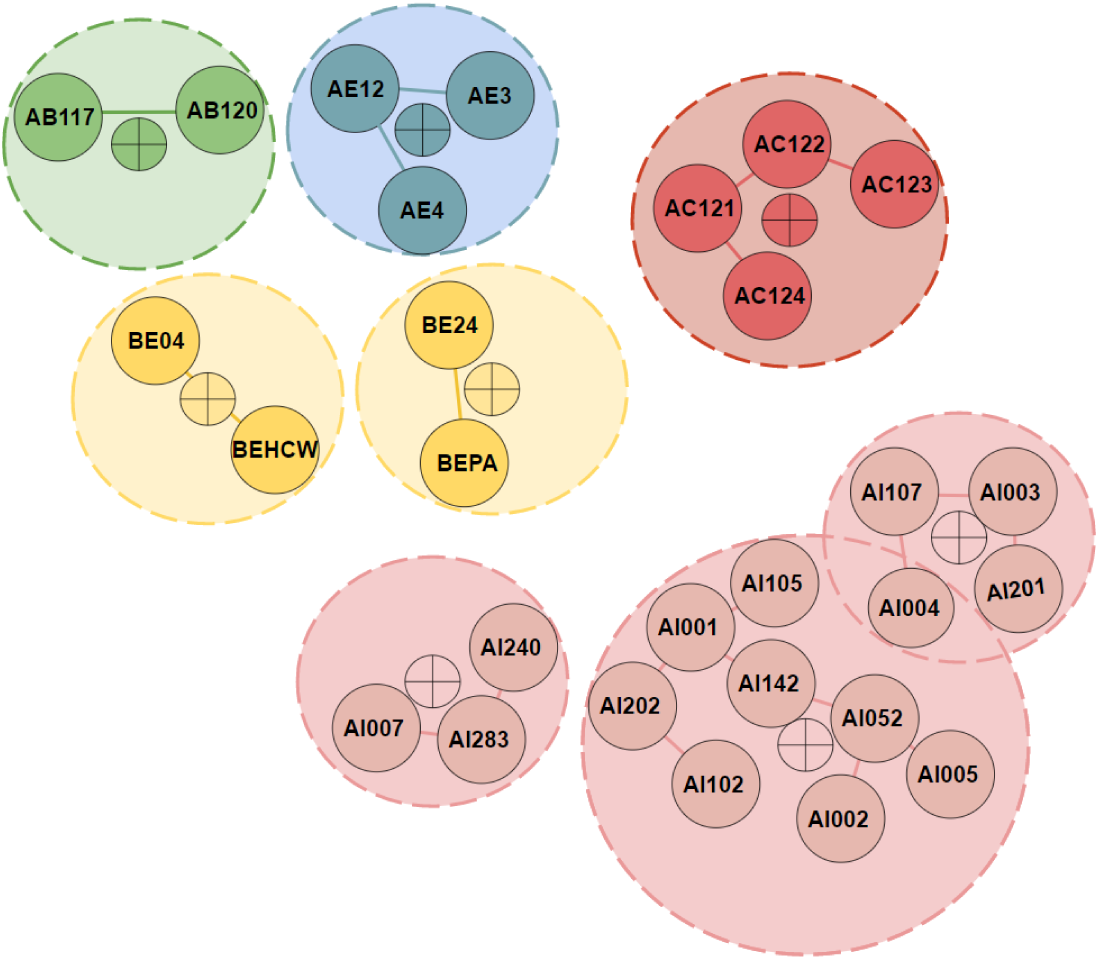
Results of hierarchical clustering. Here, clustering was applied to a set of 5 outbreaks (AB, AE, AC, BE and AI). Named circles represent samples, crossed circles represent cluster centers, obtained as arithmetic mean of corresponding clusters; hosts with the same color belong to the same outbreak. Dashed lines represent circles, where radius is equal to the distance between cluster center and farthest in-cluster host. While hierarchical clustering resulted in greater number of clusters, than viral outbreaks, adding cluster centers allowed to detect an overlap, so that the farthest in-cluster host had higher distance from the cluster center than the out-of-cluster host AI004. This way, algorithm allowed to merge 2 clusters, belonging to the same outbreak, which improved overall clustering quality.

**Fig. 6:**
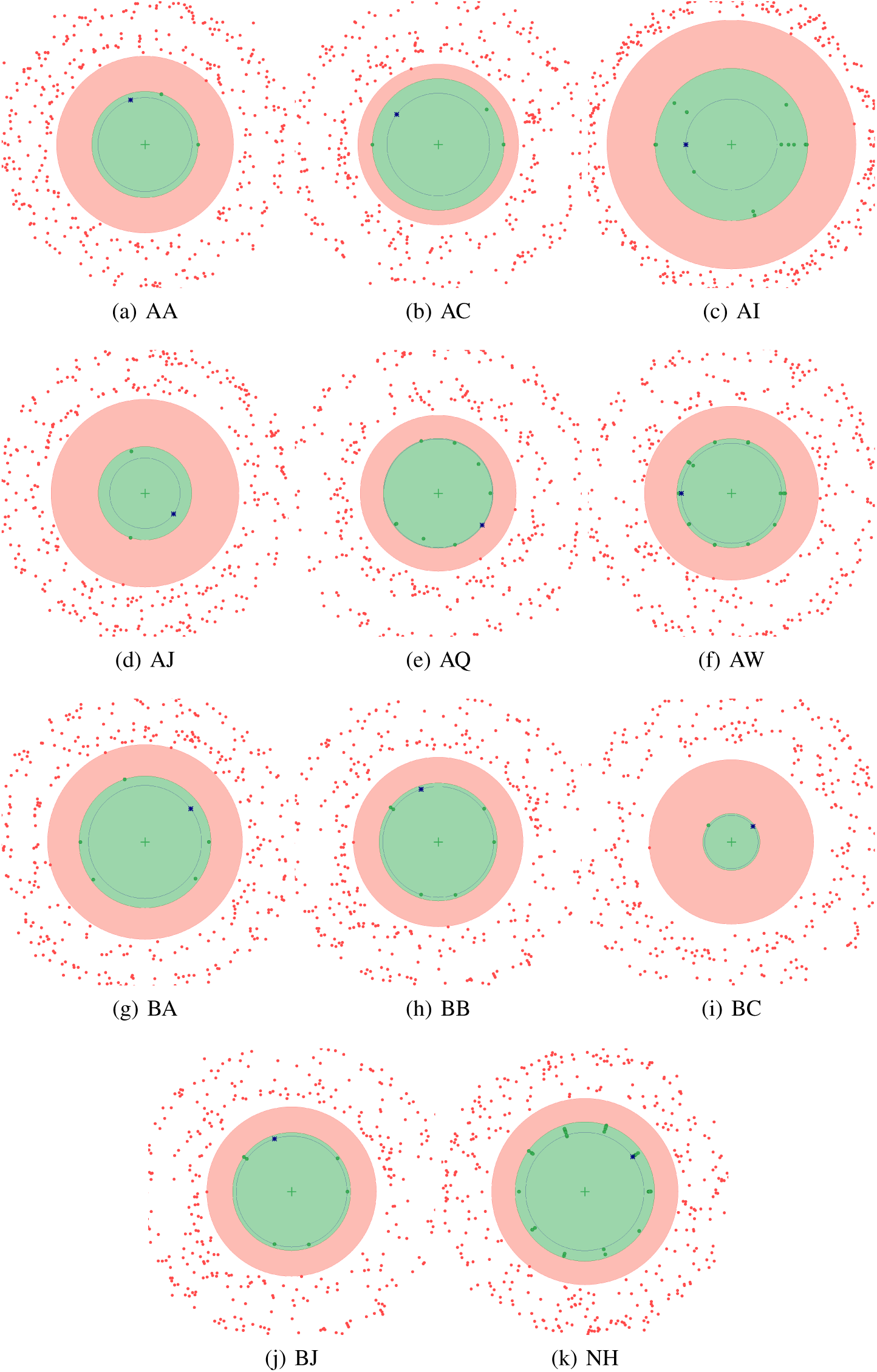
Center visualization for outbreaks with known source (k = 5). Cross in the center represents current outbreak center (represented as maximum mean); blue star represents outbreak source. Green dots represent samples, that belong to the given outbreak, red dots – unrelated samples. Distance from center represents EMD between the center and given host. For outbreak AQ, some samples are closer to the center than the source. This can be explained by the fact that in given outbreak, blood transmission was involved in virus spread.

Additionally, we find pairwise distances between all clusters, so that distance between two clusters is defined by EMD between their closes hosts, and merge clusters, that are close to each other, this way connecting clusters, that belong to the same outbreak.

### Host-to host direction inference

To infer transmission direction between a pair of infected samples, the following algorithm is applied:

1. Obtain mean between outbreak source *S* and host *X M* (*S, X*);
2. Calculate EMD between mean *M* and host *X EMD*(*M, X*);
3. Calculate EMD between mean *M* and outbreak source *S EMD*(*M, S*);
4. If *EMD*(*M, S*) *< EMD*(*M, X*), assume that infection was transmitted from *S* to *X*.

## Results

We validated our new algorithm on a publicly available dataset obtained from an epidemiological study of HCV outbreaks [2].

### Data sets

The data consists of 368 sequenced hosts where 175 of them belong to 34 annotated outbreaks. Outbreaks contain from 2 to 33 samples and 11 outbreaks have a known main spreader (Table 1). Every host sample represent an HCV intra-host population obtained with end-point limiting-dilution (EPLD). All viral sequences represent a fragment of E1/E2 genomic region of length 264bp.

**Table 1:** Outbreaks with known source

| Outbreak | AA | AC | AI | AJ | AQ | AW | BA | BB | BC | BJ | NH |
| --- | --- | --- | --- | --- | --- | --- | --- | --- | --- | --- | --- |
| # samples | 3 | 4 | 15 | 3 | 9 | 19 | 6 | 7 | 2 | 4 | 33 |

**Table 2:** Validation results. k-EMD was tested on a dataset, that includes 34 outbreaks; MinDist, ReD and VOICE were validated earlier on a smaller dataset, that didn’t include one of the outbreaks. For convenience, results for k-EMD contain 2 values – one for the smaller dataset, and one for the entire (34 outbreaks) set of hosts (values in parentheses).

| Method | k-EMD | MinDist | MinDistB | ReD | VOICE-D | VOICE-S |
| --- | --- | --- | --- | --- | --- | --- |
| **Relatedness sensitivity, %** | 90.8 (90.4) | 90 | 92.9 | 55.3 | 85.2 | 86.8 |
| **Clustering sensitivity, %** | 100 (100) | 100 | 100 | 96.3 | 98.2 | 98.2 |
| **Direction accuracy, %** | 83.9 (85.1) | N/A | N/A | 87.1 | 83.9 | 87.1 |
| **Source accuracy, %** | 80 (81.8) | 50 | 40 | 90 | 80 | 90 |

**Table 3:** Number of vertices in De Bruijn graph for corresponding k-mer sizes

| **k** | 5 | 8 | 10 |
| --- | --- | --- | --- |
| **# of vertices in De Bruijn graph** | 1024 | 28916 | 60923 |

We simulated MiSeq reads from known haplotypes using ART [9] and created mixtures using abundances from original data.

### Genetic relatedness between populations

Viral populations from two samples are genetically related if they belong to the same outbreak and unrelated, otherwise. The genetic relatedness is validated on the union of both collections containing all outbreaks and unrelated samples. There are 67528 pairs of samples, and 1007 of them are related. We used EMD as predictor for relatedness. We measured the sensitivity of our method as following. First we determining the EMD value for all unrelated pairs, the minimum value we have chosen as a threshold which prohibits false-positive relatedness detection, the pairs which have EMD below the threshold are considered as related. The sensitivity is a proportion of correctly predicted related pairs among all known related pairs. See table **??** for results.

### Outbreak prediction evaluation

We expect that all sequences in the clusters predicted by our method all belongs to the same outbreak furthermore we expect that all samples from outbreak belong to the same cluster. Similarities between true and estimated partitions were evaluated using an editing metric [3]. Given metric is defined as the minimum number of elementary operations, required to transform one partition into another, such as joining or partition of clusters[3]. Clustering sensitivity was calculated similarly to [5], so that editing distance *E* was normalized by dividing it by the number of elementary operations *N*, required to transform trivial partition into singleton sets into true partition, which is equal to *n − k*, where *n* is the number of samples and *k* is the number of true clusters[5]:

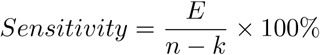

### Identification of outbreak sources

Source identification accuracy is calculated as the percentage of outbreaks with correctly predicted sources for outbreaks with known sources.

## 2 Conclusions

Extracting haplotypes by EPLD is laborious and costly procedure and that prohibits previously developed methods [5] from wide spread. On the other hand, viral samples can be easily sequenced by NGS, and that makes our novel method attractive. Furthermore, we can see that results in this article are comparable with those which were obtained using EPLD technology [5]. Moreover, our method allowed to decide whether the spreader get sequenced.

Application of molecular viral analysis to investigation of outbreaks and inference of transmission networks is a promising technique, that is available nowadays. However, it generates novel computational challenges. Given work introduced an algorithm for investigation of viral transmissions, that is based on analysis of the intra-host viral populations through k-mer decomposition. Proposed approach allows to cluster genetically related samples, infer transmission directions and predict sources of outbreaks. Validation on experimental data demonstrated that algorithm is able to reconstruct various transmission characteristics. It should be noted that even though there is still room for improvement when it comes to algorithm performance, advantage of the method is the ability to bypass cumbersome read assembly, thus eliminating the chance to introduce new errors, and saving processing time by allowing to use raw NGS reads.

## 3 Acknowledgements

A.Z. has been partially supported by NSF Grants DBI-1564899 and CCF-1619110 and NIH Grant 1R01EB025022-01. P.S. has been partially supported by NIH Grant 1R01EB025022-01 A. M. and S.K. have been partially supported by MBD.

